# Unveiling inverted D genes and D-D fusions in human antibody repertoires unlocks novel antibody diversity

**DOI:** 10.1101/2024.04.26.591287

**Authors:** Prabakaran Ponraj, Abhinav Gupta, Sambasiva P Rao, Deepak Rajpal, Maria Wendt, Yu Qiu, Partha S. Chowdhury

**Affiliations:** Large Molecules Research, Sanofi, Cambridge, MA, USA; Translational Science, Sanofi, Cambridge, MA, USA; Takeda, Development, Lexington, MA, USA; Johnson & Johnson R&D Center, Spring House, PA, USA

## Abstract

Antibodies, fundamental to immune defense, derive their diversity primarily from the intricate rearrangement of variable (V), diversity (D), and joining (J) gene segments. Traditionally, D genes in the forward (5’-3’) direction contribute to this diversity by rearranging with V and J segments. However, the existence and significance of inverted D genes (InvDs), which are D genes oriented in the inverted (3’-5’) direction, were previously obscured by limitations in data and detection methods. Here, we carried out a comprehensive analysis of a large-scale public next-generation sequencing (NGS) dataset encompassing antibody repertoires from 13 healthy donors using a novel immunoinformatics workflow. Our analysis, for the first time, uncovers the existence of all 25 unique InvDs across all three reading frames within human antibody repertoires, including both naïve and memory B cells. This finding challenges previous assumptions, revealing the extensive presence of InvDs and identifying a broad spectrum of D-D fusions, especially those involving InvDs. Notably, InvDs enrich for unique amino acids such as histidine, proline, and lysine, not commonly found in forward D genes, and exhibit reduced use of certain negatively charged and bulky amino acids, including aspartate, tryptophan, and methionine. The unique amino acid profile of InvDs discloses new diversity and functionality in the human antibody repertoire, evidenced by over two dozen documented antibodies featuring InvDs, targeting a wide array of antigens. By opening exciting avenues for immunogenetics research, including new chromatin compaction models, innovative antibody libraries, and advancements in antibody engineering, these findings hold promise for the development of novel therapeutics and vaccines.

## Introduction

The remarkable diversity of antibodies, crucial for neutralizing a vast array of pathogens, arises from a unique genetic process involving the recombination of variable (V), diversity (D), and joining (J) gene segments in B cells (*1–3*). D genes are instrumental in sculpting the major antigen-binding site, the complementarity-determining region 3 of the heavy chain (CDR-H3), via their recombination with V and J segments(*4, 5*). This process is further refined by the insertion of random P- and N-nucleotides at the VD and DJ junctions, along with meticulous trimming at the ends of D genes, enhancing the complexity of CDR-H3. The recombination is governed by the 12/23 rule, where V and J segments with 23 base pair (bp) spacers are paired with D segments flanked by 12 bp spacers in their recombination signal sequences (RSS). Given that D gene segments have RSS at both 5’ and 3’ ends, they can potentially rearrange in both direct and inverted orientations and are theoretically capable of encoding in all three reading frames (RFs)(*6*). D genes also have the potential for further diversity through tandem D-D fusions(*7, 8*). Previous studies, using limited in vitro and transgenic models, identified only a few inverted D genes (InvDs) and D-D fusions(*9–13*). Despite the use of Sanger and 454 sequencing for antibody repertoire analysis, subsequent studies continue to find that InvDs in the expressed human antibody repertoire remain elusive(*14, 15*). This is likely due to limitations in the sequencing depth or detection methods employed in earlier studies.

The rapidly evolving fields of B cell biology and antibody therapeutics research face a critical unanswered question: the true extent of InvDs within the human antibody repertoire and their impact on antibody diversity and function. To address this, we analyzed publicly available antibody repertoire sequencing data from 13 healthy individuals. For 10 donors, the data comprised nearly 3 billion annotated V_H_ sequences, representing over 106.5 million unique antibodies with available in-frame V_H_ and CDR-H3s, alongside IGHV, IGHD, and IGHJ gene families(*16*). For the remaining 3 donors, data provided nearly 30 million unique CDR-H3s of antibodies, including both naïve and memory B cells(*17*). This comprehensive analysis of over 157 million unique antibody sequences revealed a surprising prevalence of InvDs from all D gene families across all three reading frames, not only in naive B cells but also in memory B cells. A two-step approach identified potential InvDs: first, a nucleic acid-based BLAST search based on sequence similarity and alignment length to known InvD germline sequences, considering InvD length during the search. Second, machine learning-based clustering and visual inspection of the amino acid sequences of CDR-H3s confirmed the InvDs utilizing all three reading frames. Notably, our findings not only unveil the existence of InvDs but also highlight their far greater role in shaping antibody diversity than previously understood. Additionally, we explored the involvement of InvDs in D-D fusions, a mechanism that contributes even more to the remarkable diversity of the antibody repertoire (Fig. 1). Distinct amino acid trends associated with InvDs across all reading frames, particularly the presence of histidine-rich and proline-rich stretches uncharacteristic of D genes in direct orientation, might influence antibody function. In support of this, we identified more than two dozen human antibodies documented in the literature and patents that contain InvDs within their antigen-binding regions (CDR-H3), potentially targeting a diverse array of antigens. This finding underlines the practical importance of InvDs for developing effective therapeutics. Overall, these results hold significant promise for advancing antibody research by opening new avenues, particularly relevant for therapeutic development. By highlighting the previously unrecognized role of InvDs, we pave the way for more comprehensive investigations into antibody diversity, ultimately leading to a broader spectrum of treatments for various pathogens and diseases.

**Fig. 1.**
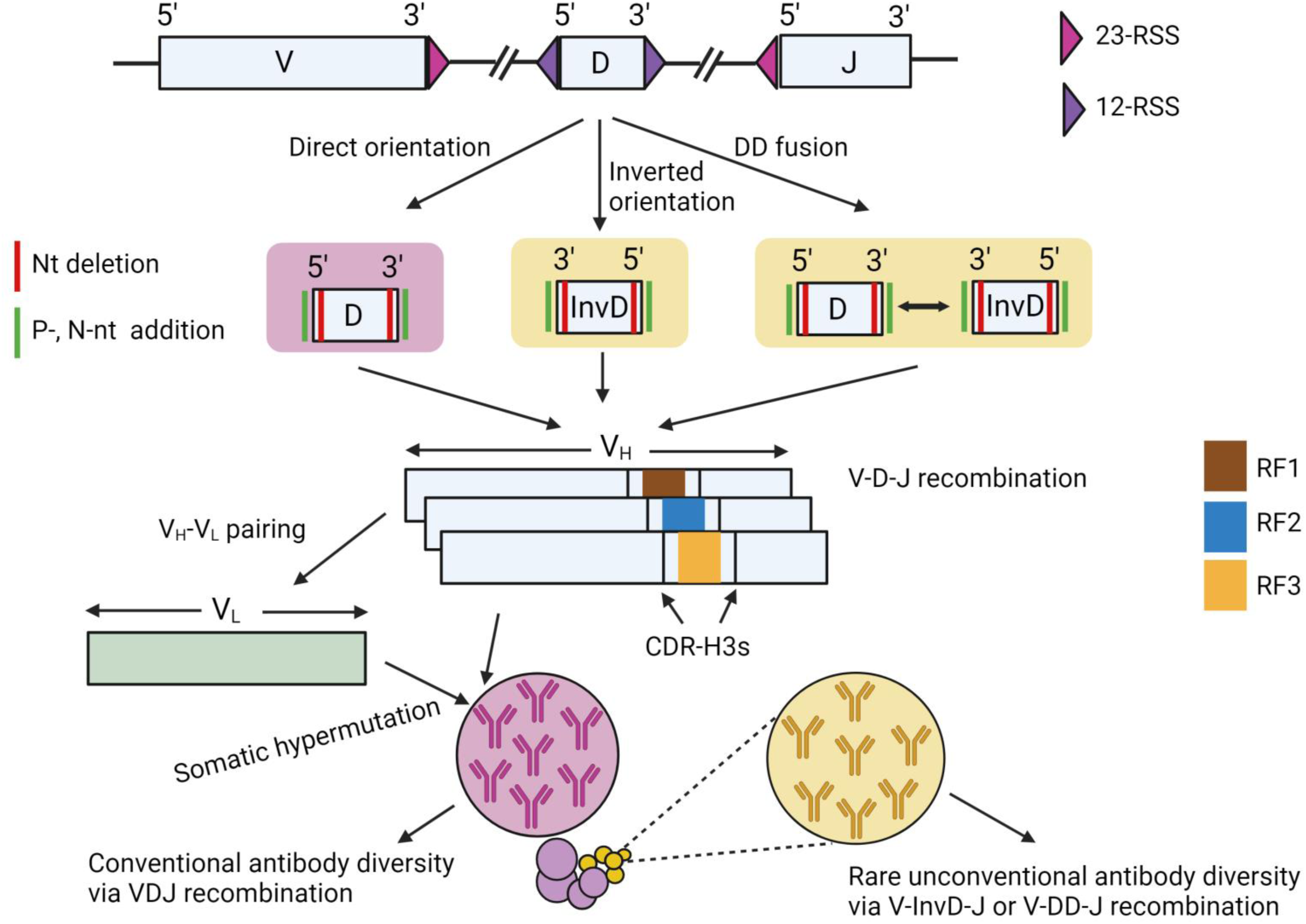
D genes as conventional and unconventional diversity sources in human antibody repertoires. D genes facilitate VDJ recombination through 12-bp and 23-bp recombination signal sequences (RSS), following the 12/23 rule. They recombine in direct (D) or inverted (InvD) orientations with junctional modifications like nucleotide (nt) deletion and P, N-nt addition, utilizing any of the three reading frames (RF1, 2, and 3). Notably, D genes also undergo DD fusion, combining D and InvD. This versatility in D genes, creating CDR-H3s in the heavy chain (V_H_) and influencing light chain (V_L_) pairing and somatic hypermutation, contributes to the vast diversity of human antibody repertoires. The rare, unconventional diversity from InvDs and DD fusions enhances our understanding of immune complexity, revealing potentially novel and previously unknown paratope space.

## Results

### Length-categorized stringent criteria applied to BLASTn results reveal all InvDs in naïve and memory antibodies across donors, demonstrating their role in antibody diversity

The human antibody repertoire contains 27 human D genes, and 25 of them are unique (Table S1)(*18*). These genes, whose lengths range from 11 to 37 nucleotides (nt), are prone to truncation, extensive junctional modifications, and potential somatic hypermutation (SMH), complicating their identification compared to V and J genes. Previous studies, including extensive work by Collins and coworkers on D gene annotation have utilized varying nucleotide length alignments (approximately 6 to 11 nt) and sequence identities with 0-2 mismatches to corresponding germline D genes(*19–21*). Several tools such as SODA2, IgBLAST, iHMMune-align, and BRILIA, all of which have contributed to the identification of D genes(*22–25*). Nonetheless, InvD identification efforts frequently resulted in improbably short InvD alignments with germline sequences, leading to ambiguous identifications or frequencies of occurrence that were much lower than the detection methods’ precision threshold, thus preventing the identification of InvDs(*15, 25*).

To overcome these limitations of short and unreliable InvD alignments, we developed a method that categorizes germline InvDs based on length for BLASTn queries: short (11-18 nt), medium (19-28 nt), and long (31 and 37 nt). We first implemented rigorous nt-based BLASTn searches followed by filtering based on tailored length and identity thresholds for each InvD category and specific cases (Fig. 2). Through alignments allowing for some mismatches, we observed that a few of the InvD sequences (e.g., InvD6-6, InvD6-25, and InvD6-13) encode the ‘CX4C’ amino acid motif (C = cystine; X = any amino acid), which is also encoded by D2-2 in the standard orientation (Table S1). To improve reliability for such cases, we excluded InvDs with the “CX4C” amino acid motif in their CDR-H3s. This approach yielded a substantial number of InvDs: 32,236 short, 44,121 medium, and 37,912 long, all with high sequence similarity and alignment correspondence to their germline counterparts (Fig. S1, Table S2, S3 and S4), demonstrating the effectiveness of our methodology.

**Fig. 2.**
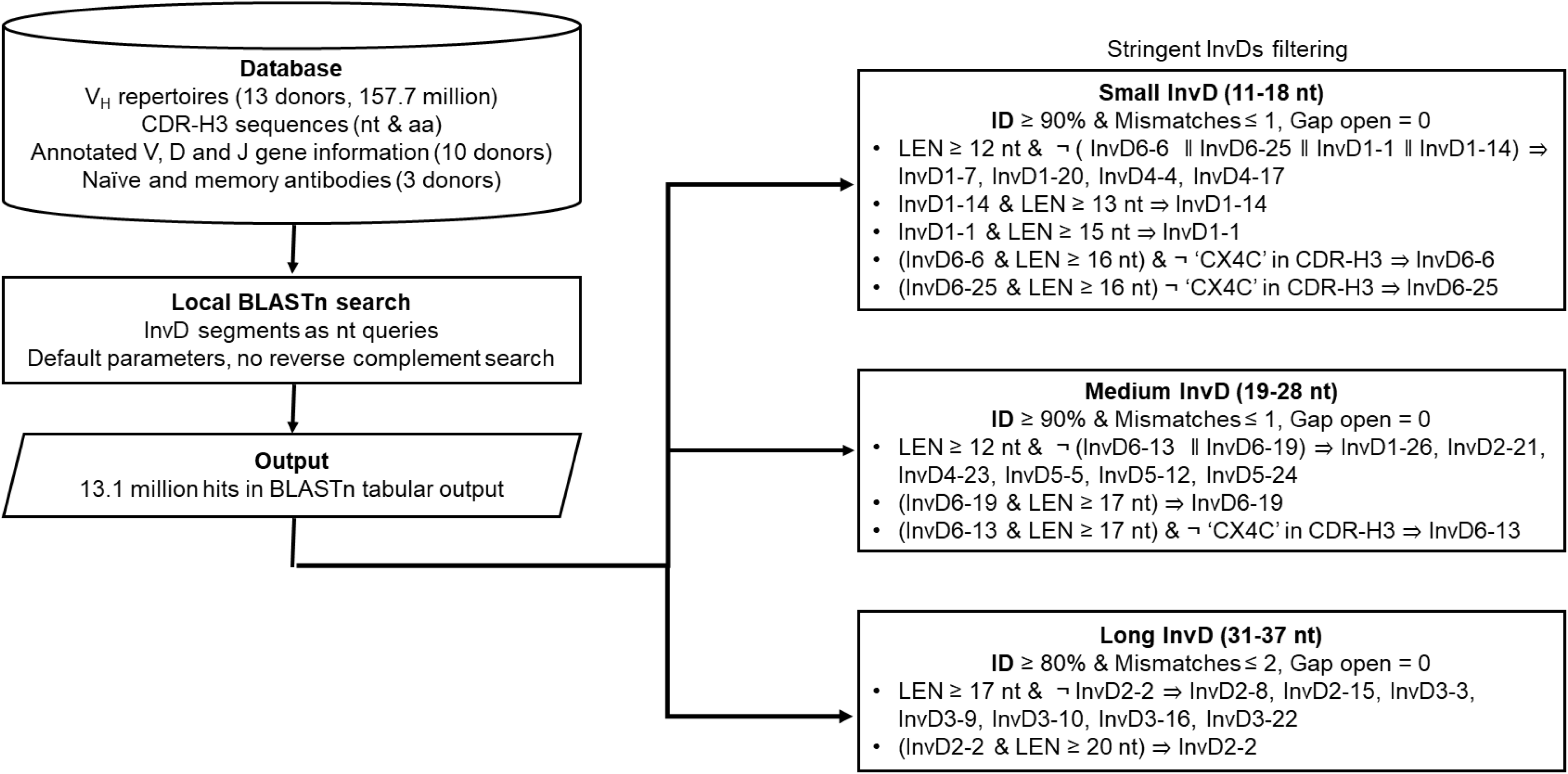
Workflow for identifying Inv-Ds in human antibody repertoires. This methodology highlights the process of detecting InvDs within antibody heavy chain variable domain (V_H_) repertoires from 13 donors. The process begins by compiling CDR-H3 sequences in both nucleotide (nt) and amino acid (aa) forms. Next, a local BLASTn search is conducted against known InvDs. After classifying InvDs by length (short, medium, and long), the resulting output undergoes stringent filtering using SAS JMP Query Builder, applying criteria based on sequence identity and alignment length. Search refinement employs nt and aa level criteria, using logical operators (“&” for “AND” and “¬” for “NOT”). Additionally, the exclusion of the “CX4C” motif helps differentiate overlapping InvDs and direct orientation D gene segments (refer to the text for details). The final selection of InvDs meeting these criteria ensures reasonably accurate InvD identification with significant sequence similarity and alignment correspondence to germline InvDs (Fig. S1).

The violin plots depicting the InvDs associated with antibody sequences from 10 donors showed distinct distributions across the specified length categories (Fig. 3a). Noticeably, memory cell derived antibodies, alongside naïve ones, displayed considerable diversity, suggesting functional significance of InvDs in antigen recognition and immune response mechanism (Fig. 3b). Further, a detailed analysis of 13 donors revealed the usage of individual InvD gene families within naïve and memory antibodies across different length categories (Fig. S2). Thus, our analysis revealed widespread usage of all InvD gene families across different length categories within healthy human antibody repertoires, contradicting previous reports of limited or no InvD presence(*14, 15*). The presence of InvDs in both naïve and memory antibodies is particularly remarkable, suggesting a potential role for InvDs in shaping antibody diversity across different stages of the immune response.

**Fig. 3.**
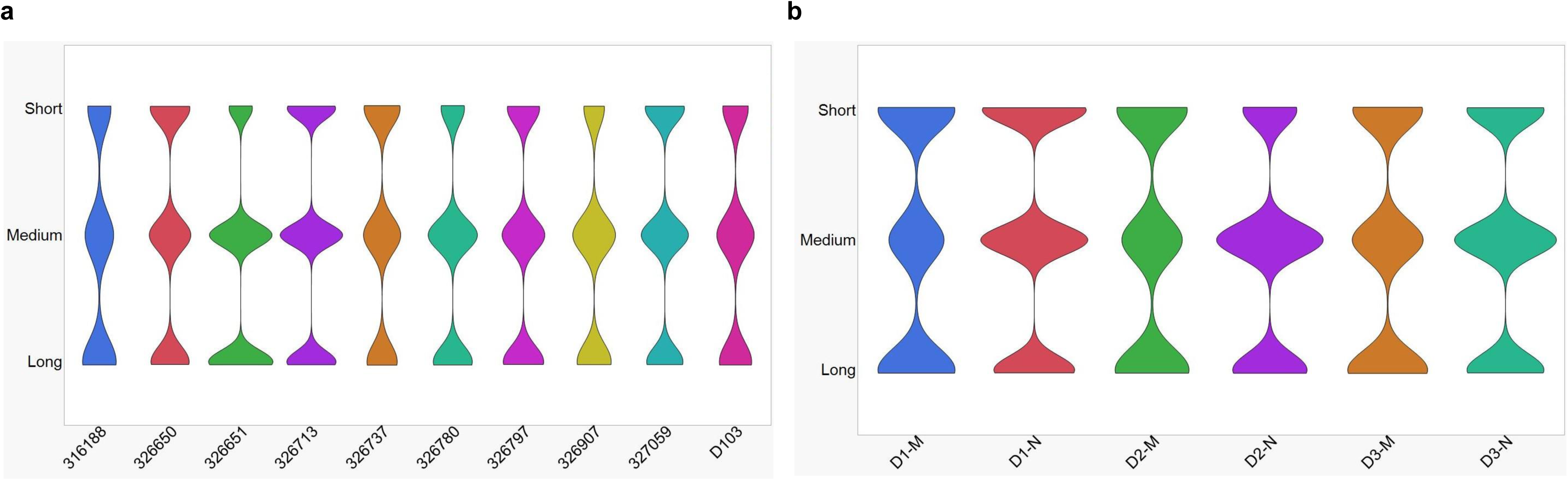
Prevalence and distribution of InvDs in human antibody repertoires. **a**, Violin plots illustrate the existence and distribution of on InvDs within the expressed antibody repertoires of 10 donors. These InvDs are categorized by length: short (11-18 nt), medium (19-28 nt), and long (31-37 nt). The X-axis distinguishes individual donors through color coding, while the Y-axis in the violin plots represents the relative abundance of InvDs. **b**, For additional 3 donors, violin plots further dissect the prevalence of InvDs within naïve (N) and memory (M) antibodies. This suggests that InvDs may play a significant role in the maturation and antigen recognition capabilities of the immune system. The breadth of each violin plot serves as an indicator of the variety and abundance of InvDs within the CDR-H3 regions of heavy chain antibody repertoires. See Fig. S2 for analysis of InvD gene families across 13 donors with length categories and for naive vs memory B cells for 3 donors.

### InvDs promote antibody diversity by utilizing all three reading frames, enabling unique paratope diversity

After we confirmed the prevalence of all InvDs in the expressed CDR-H3s of antibody repertoires in normal humans, we then turned our attention to the analysis of inverted reading frames. D genes can potentially be used across all three reading frames (RF1, RF2 and RF3) in both direct and inverted orientations - RF1 initiates with the first codon of the D coding region, while RF2 and RF3 are derived from one and two nucleotide shifts, respectively. This allows three different amino acid sequence motifs encoded by each of the regular D and InvD segments (Table S1). To explore the usage of inverted reading frames, we performed unsupervised clustering of InvDs-associated CDR-H3s, as described in Fig. S3, using the ESM2 embedding(*26*). Using this methodology, we were able to show for the selected InvDs - InvD6-25, InvD6-13, and InvD2-2 representing small, medium, and long lengths, respectively, illustrating distinct clusters with the three reading frames (RF1, RF2, RF3), post mapping of germline-encoded RFs (Fig. 4a-c). Extending this analysis to all InvDs, we found that distinct clusters indicating three RFs usage for certain InvDs, while other patterns suggest mixed RFs or non-clustered arrangements (Fig. S4), possibly due to extrinsic factors, data constraints, or the lack of model’s attention to InvDs. Thus, our analysis revealed that all InvDs utilize all three reading frames, RF1, RF2, and RF3, even in the presence of stop codons at some of the RF2s. These stop codons may be eliminated during V(D)J recombination through nucleotide addition or deletion, potentially allowing the corresponding InvDs to contribute functional antibody sequences(*27*).

**Fig. 4.**
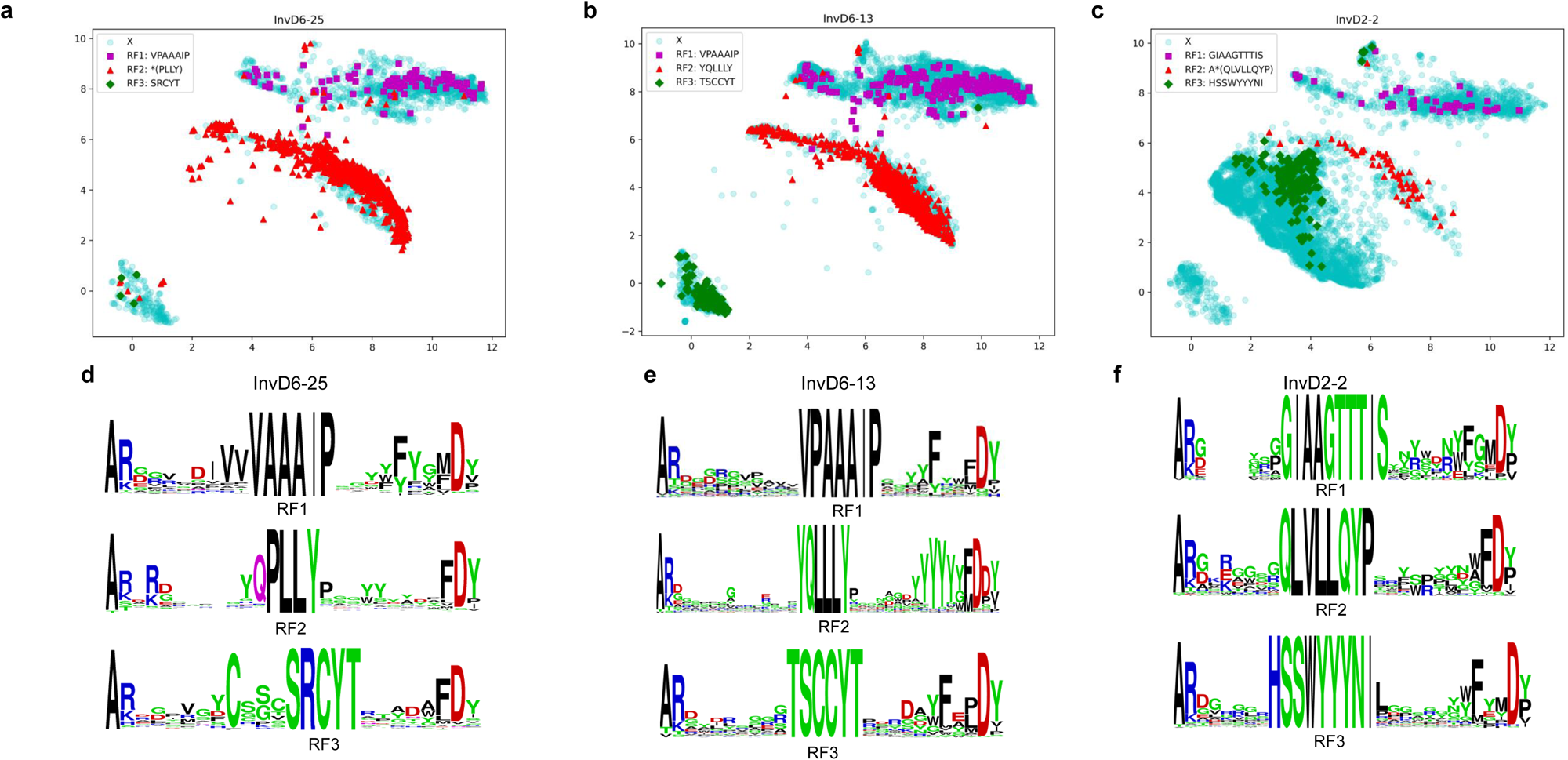
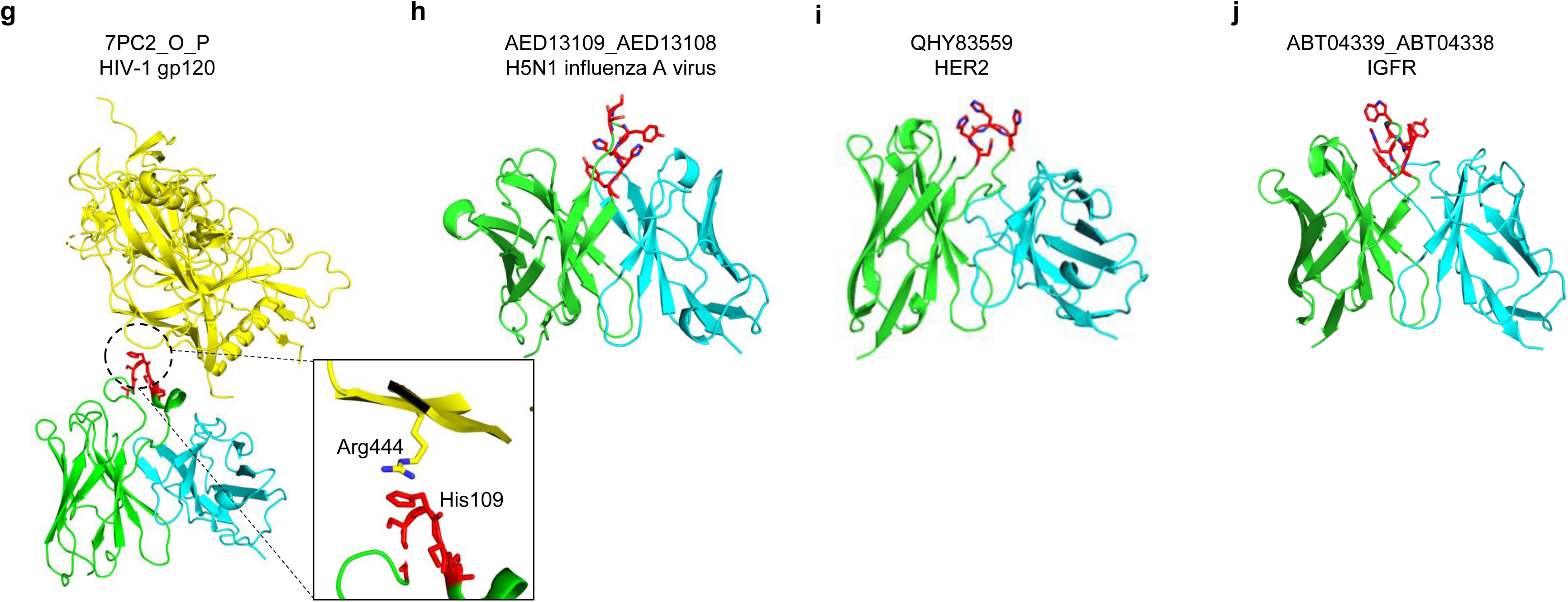
InvDs Contribute to CDR-H3 diversity by employing all three reading frames (RF1, RF2, and RF3) and functional antibodies. **a-c**, Zero-shot clustering of CDR-H3s using mean embeddings from the ESM2 model (methodology in Fig. S3). The selected InvDs—InvD6-25, InvD6-13, and InvD2-2—represent small, medium, and long InvDs, respectively. Inverted reading frames (RF1, RF2, and RF3) are mapped to each InvD-associated CDR-H3 sequence based on a comparison with its germline amino acid sequence. If the full sequence does not match, the motif given within the brackets is used for mapping. The UMAP projections are color-coded according to the assigned reading frames: purple-colored squares for RF1, red-colored triangles for RF2, green-colored diamonds for RF3, and unassigned CDR-H3s are denoted by a cyan-colored circles. Interestingly, ESM-2 clusters CDR-H3s according to RFs even though such information was never used in its training. **d-f,** Sequence logos illustrate the utilization of all three reading frames (RF, RF2 and RF3) by these InvDs, highlighting the remarkable diversity generated within CDR-H3s. Notably, sequences originating from RF2, despite harboring germline-encoded stop codons, actively contribute to functional CDR-H3s. This observation suggests a fascinating process of excision during recombination, allowing these sequences to play a crucial role in antibody diversity. **g-j,** Human antibodies harboring InvDs within their CDR-H3s, identified from the Patent and Literature Antibody Database (PLAbDab), specifically target HIV-1, H5N1 influenza virus, HER2, and IGFR. Shown are the 3D structural models, except for one (PDB 7PC2) where the complex crystal structure with the antigen, HIV-1 gp120, is available and shown in yellow. The heavy chain is depicted in green, the light chain in cyan, and germline residues matching the InvDs within the CDR-H3 regions are highlighted in red. The corresponding PLAbDab IDs and target names are provided for each antibody model. See Fig. S8 for more examples and Supplementary Table 6 for details.

To further investigate the utilization of all three RFs in InvD-associated CDR-H3s, we employed sequence logos(*28*). We selected CDR-H3 sequences containing germline-encoded sequence motifs specific to each RF, namely, RF1, RF2, and RF3, from representative InvDs of varying lengths: InvD6-25 (short), InvD6-13 (medium), and InvD2-2 (long) (Fig. 4d-f). These amino acid sequence logos clearly demonstrated the presence of all three RFs within these InvDs, highlighting the remarkable diversity generated in CDR-H3s. Furthermore, we generated sequence logos for all InvDs, encompassing a broader range of lengths and amino acid properties, but still focusing on sequences with germline motifs corresponding to all three RFs. These logos confirmed the utilization of all three reading frames, RF1, RF2, and RF3, in generating diversity within CDR-H3s across the entire InvD population (Fig. S5).

Previous studies in mice suggested a preference for a single reading frame in D genes, often encoding tyrosine and glycine. Other reading frames, enriched in hydrophobic and charged amino acids were thought to have a negative impact on B cell development and antibody function(*29–32*). Similar patterns were reported in human studies, where most D genes were found to favor a single reading frame, and the use of inverted reading frames was uncommon(*14, 15*). However, our analysis presents a new perspective on the human antibody repertoire, challenging these established views. We observed that all InvDs, present in both naive and memory states, utilize all three reading frames. This remarkable adaptability leads to distinct amino acid usage patterns compared to uninverted D genes. Notably, histidine and proline are significantly enriched in the InvD-derived germline sequences (4.7% and 8.3%, respectively) (Table S5), potentially influencing the composition of certain germline InvD-derived CDR-H3s (Fig. S5). This increased diversity in CDR-H3s, driven by InvD reading frame usage, might contribute to enhanced antibody specificity and precision in recognizing a broad range of antigens.

### InvDs contribute to extensive VDJ diversity and enhance specificity towards a wide range of antigens

To investigate the effect of InvDs on antibody repertoire diversity via V/J gene usage and V(D)J recombination, we initially examined the impact of InvD length. Surprisingly, short, medium, and long InvDs exhibited no significant differences in V and J gene family usage (Fig. S6a, b), suggesting minimal influence of InvD length on partner selection during V(D)J recombination. While InvDs come in various lengths, their frequencies also differ within each category (Fig. S6c-e). This diversity within InvDs likely contributes to overall antibody repertoire variation. Further, VDJ diversity plots for different InvD lengths (Fig. S7a-c) reveal a rich landscape of recombination events between V, D, and J gene segments, with the color gradient emphasizing the extensive diversity across all categories. Collectively, although InvDs do not directly determine V and J gene family selection, their intrinsic length variations and distinctive reading frame usage patterns play a significant role in shaping antibody repertoire diversity. This enables a remarkable spectrum of VDJ recombinations, potentially resulting in a more functional and diverse antibody repertoire.

To further substantiate the potential role of InvDs in shaping antibody specificity, we investigated human antibodies harboring InvDs within their CDR-H3s. These antibodies were identified from the Patent and Literature Antibody Database (PLAbDab)(*33*). Analysis of their 3D structural models, as available in the database, revealed the presence of InvDs highlighted in red within the CDR-H3s (Fig. 4g-j, Fig. S8 for more examples). Notably, a complex crystal structure of an anti-HIV antibody (PDB 7PC2) bound to its antigen, HIV-1 gp120 (visualized in yellow in Fig. 4g), provided a unique opportunity to explore potential interactions. This detailed structural information revealed a cation-pi interaction between a histidine residue within the germline-encoded portion of the InvD6-19 (highlighted in red in Fig. 4g) and an arginine residue from the HIV-1 gp120 epitope. This interaction (Fig. 4g) exemplifies how InvDs can contribute to antigen recognition through novel biochemical mechanisms. Furthermore, analysis of 24 unique antibodies targeting a diverse range of infectious agents (HIV-1, SARS-CoV, influenza virus, Ebola virus, and anthrax) and human antigens (IGFR, activin A, HER2, CD3, and CD74) revealed a remarkable structural and genetic diversity (Fig. S8, Table S6). These InvD-containing functional antibodies originated from various V genes (IGHV, IGKV, and IGLV germline genes) and utilized diverse InvD genes with different reading frames (Table S6). Overall, the specific targeting of diverse antigens by antibodies harboring InvDs suggests a potentially significant contribution of InvD reading frame usage to a broader and more precise antibody response.

### InvDs amplify antibody diversity complexity through D-D fusions, adding paratope diversity with long CDR-H3s

As an additional methodology to identify InvDs and explore potential D-D fusion events involving regular D segments and InvDs, we employed the IgScout algorithm(*34*). IgScout analysis identified the frequencies of all InvDs across 13 donors (Table S7). This analysis revealed a significant enrichment of InvDs in both naive and memory B cell repertoires, with an average frequency of 0.44% in naive B cells, increasing notably to 1.23% in memory B cells (Table S7). This memory B cell enrichment aligns with our observations of a broader distribution of InvD gene families within the memory B cell population compared to naive B cells (Fig. S2d-f). Furthermore, the IgScout analysis of D-D fusions revealed a remarkable diversity in these events, incorporating direct Ds, InvDs, and combinations thereof, labeled as D-D, D-InvD, InvD-D, and InvD-InvD (Figs. S9, S10 and Table S8). These D-D fusions associated with diverse V and J genes, contributing to longer CDR-H3s up to 40 amino acid length (Table S9). These results provide compelling evidence for the extensive involvement of InvDs in D-D fusions within the human antibody repertoire, further emphasizing their significant contribution to the generation of antibody diversity.

## DICUSSION

Antibodies have become the leading class of therapeutic drugs, uniquely binding to a broad range of targets with diverse affinities, while meeting high safety and functional standards(*35*). Central to their efficacy, antibody diversity, particularly the CDR-H3, is essential to defining their binding specificity and functional properties(*4*). In this study, we analyzed large-scale antibody sequence datasets, demonstrating that InvDs significantly contribute to the unique diversity of the human antibody repertoire. This has potential implications for antibody specificity and immune responses, which are crucial for the development of antibody-based therapeutics and vaccines.

Previous studies have reported limited or no InvD presence and infrequent D-D fusion events, considering them rare occurrences and their utilization of inverted reading frames even more exceptional (*9, 10, 14, 15, 36*). Particularly, the few InvDs associated antibodies previously identified were often associated with autoimmune and other disease states(*37–40*). In contrast, our extensive analysis revealed all 25 unique InvDs were present across all three reading frames within the antibody repertoires of healthy individuals. This widespread presence across reading frames highlights the potential of InvDs to significantly diversify antibody repertoires. Furthermore, we observed a significant enrichment of both InvDs and D-D fusions involving InvDs in both naive and memory B cells. This challenges the prevailing notion of InvD rarity and limited functionality(*5, 15, 41, 42*). Validating InvD function in antibodies, we analyzed human antibodies from the PLAbDab harboring InvDs in their CDR-H3s (*33*). These antibodies target diverse antigens, including viral proteins (HIV, SARS-CoV, influenza, Ebola) and human proteins (HER2, CD3, IGFR, activin A, CD74). Notably, D-D fusion antibodies, especially those with long CDR-H3s involving regular D and InvDs, may excel at binding elusive targets like cryptic epitopes, ion channels, and GPCRs. These findings suggest that InvDs play a crucial role in shaping the human antibody repertoire, potentially contributing to a broader range of immune responses, including the generation of long-lived memory B cells essential for pathogen defense.

Traditionally, limited knowledge of D genes, their reading frames, and how they play a role in amino acid sequences undergoing positive and negative selection pressures raised questions about the extent of the D gene’s contribution to the diversity of the human antibody repertoire(*5, 41*). Our analysis confirmed, in humans, the usage of all 25 unique InvD gene families across all three reading frames with functional potential, opening a previously untapped source of diversity, potentially enabling the production of a broader array of antibodies with unique specificities. For example, unlike conventional D segments, InvDs encode more histidine and proline residues, critical for antibody-antigen interactions(*43*). This allows InvD-derived antibodies, particularly those rich in these residues, to achieve high affinity early on, potentially bypassing extensive maturation. Histidine in InvDs also might enable pH-dependent binding, offering targeted action in specific environments(*44*). Additionally, InvD-derived antibodies enriched with such histidine residues may exhibit an extended half-life, leading to longer-lasting therapeutic effects(*45*). These unique properties hold promise for developing novel and more effective antibody-based therapies(*46*).

In InvDs, nine non-canonical germline-encoded single cysteines are present through their RF3. If these single cysteines are not exposed to the surface, they could contribute to stability through hydrogen bonding(*47, 48*). Additionally, a CC motif is found in InvD6-13 through the RF3. This CC motif may play an important role, like vicinal disulfides in proteins, including carbohydrate binding(*49*). Interestingly, several long CDR-H3s with D-D fusions have germline-encoded non-canonical double cysteines. These double cysteines involve any of the regular D2 gene families (D2-2, D2-8, D15, and D-21) through the RF2. These long CDR-H3s have the potential to form intra-CDR-H3 disulfide bonds, which could rigidify their extended loops and bind to diverse antigens(*50*).

Furthermore, the potential of InvDs to diversify the antibody memory repertoire opens exciting avenues for vaccine development, particularly for previously difficult targets. InvD-associated antibodies, especially from underutilized RF1, hold promise due to their enrichment in positive charges compared to those containing regular D segments(*51*). This may be advantageous for vaccines like HIV-1, where the virus has a negatively charged envelope. By designing B-cell activating immunogens that specifically target germline antibodies enriched for InvDs, one could leverage these unique properties to significantly enhance vaccine efficacy against elusive pathogens. This approach has the potential to elicit more robust and broadly neutralizing immune responses(*52*).

In conclusion, our findings suggest that D genes go beyond their known role in conventional antibody diversity. They exhibit a remarkable ability to undergo inversion and fusion, with significant implications for antibody therapeutics research and vaccine design. This phenomenon mirrors how gene inversion and duplication contribute to broader genomic diversification and the emergence of novel functions. Finally, it is important to acknowledge that our stringent search conditions could have introduced bias or led to an underestimation of the actual prevalence of InvDs, especially those with significant mutations or truncations. Nonetheless, our large dataset-driven analysis has yielded crucial results that will undoubtedly prompt future experimental work and the creation of computational tools and functional studies. The observation of extensive InvDs and D-D fusions necessitates an understanding at the chromatin level and the use of compaction models to elucidate the creation of complex, unconventional diversity(*53–55*). Further efforts will be dedicated to investigating novel human antibodies generated through InvD and D-D fusions, to develop innovative therapeutic strategies.

## Acknowledgements

We thank Melody Shahsavarian and Adrian Carr from the Large Molecules Research for their critical reading of the manuscript and suggestions on the figures, respectively. We also thank Ally Hatton for her business and legal support, and Dongyu Liu from Translational Science for technical assistance.

## Author contributions

Conceived and designed the study: P.P., P.S.C. Performed analysis: P.P., D.L., A.G. Supervised the research: P.S.C., M.W., D.R., F.R., S.K., Y.Q. Wrote the paper: P.P. Edited and approved the final manuscript: All authors.

## METHODS

Next-generation sequence datasets of human antibody V_H_ repertoires from 10 and 3 individuals were compiled from the previously published studies(*16, 17*). For the dataset of 10 donors, all unique in-frame V_H_ amino acid sequences with annotated IGHV, IGHD and IGHJ germline gene families, CDR-H3s were extracted using UNIX scripts. All IGHV, IGHD and IGHJ germline gene families in the dataset A were originally annotated by the authors using Abstar (version 0.3.3). It should be noted that annotated CSV and JSON files as downloaded from database contained the consensus sequences from nearly 3 billion V_H_s. However, most of the clusters contained only a single sequence and many of the consensus sequences could be identified in the raw datasets. For the dataset of 3 donors, productive and unique V_H_ sequences were extracted from both memory and naive B cell receptors.

The CDR-H3s of both nucleic acid (nt) and amino acid (aa) sequences were retrieved from the databases and stored along with the available annotated information – V, D and J for 10 donors and naïve and memory information for 3 donors, in JMP tables. The FASTA files containing nt sequences tagged with annotated information and aa sequences were prepared for 13 donors using shell scripts. The query nt sequences containing all 25 unique InvDs (Table S1) were prepared from the IMGT database (*56*). The nt-based local BLASTn search was performed by scritping(*23*). Custom workflows for identifying InvDs, after separating them into small (11-18 nt), medium (19–28) and long (31–37), with stringent matching conditions were applied using JMP Query Builder (Fig. 2). These processes produced hits - 32,236 for small, 44,121 for medium and 37912 for long categories, with potentially InvD-associated CDR-H3s (Supplementary Tables 3, 4 and 5) which were used for the analysis InvDs and RFs. The nt sequences along with lengths corresponding to Ds and InvDs with three RFs, aa sequences, (Table S1), and aa composition for Ds and InvDs (Table S2) were complied. All data and statistical analysis and plots were made with JMP Pro 17.2.0. Zero-shot clustering of all InvDs containing CDR-H3s was performed using mean ESM2 embeddings (Fig. S3). IgScout program(*34*) was used with InvDs nt sequences as queries for identifying the usages of InvDs and D-D fusion (Table S6 and 7). The PLAbDab database of human antibody sequences was scanned using local BLAST alignment to identify antibodies with sequences matching known InvD germline sequences(*33*).

No statistical methods were used to predetermine sample size. The experiments were not randomized, and investigators were not blinded to allocation during experiments and outcome assessment.

## Data availability

All data analyzed are included in this Analysis and its Supplementary Information.

## SUPPLEMENTARY MATERIALS

**Fig. S1.**
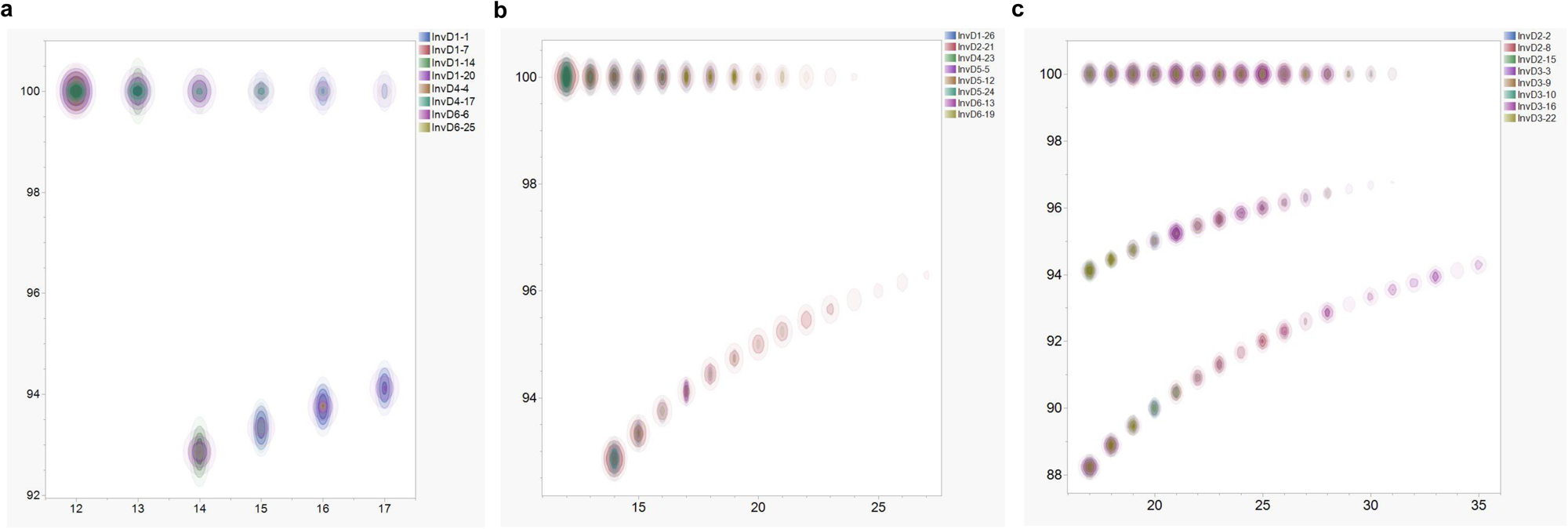
Contour plots representing the distributions of short (11-18 nt), medium (19 – 28 nt) and long (31-37 nt) InvDs identified within human antibody repertoire datasets. These plots illustrate the stringent filtering criteria applied to identify InvDs among 13 donors’ repertoires. Axes indicate nucleotide (nt) lengths (X-axis) and sequence identity percentages (Y-axis) relative to their corresponding D genes in inverted orientation, following the parameters outlined in Fig. 1. **a**, Identification of short InvDs, highlighting the high sequence identity and length, with InvD7-27 addressed separately due to its minimal length. **b**, Reliable identification pattern for medium InvDs, denoting a robust match in sequence identity and length. **c**, The distribution of long InvDs, showing the prevalence and range of long InvDs within specified high-confidence identity and length criteria.

**Fig. S2.**
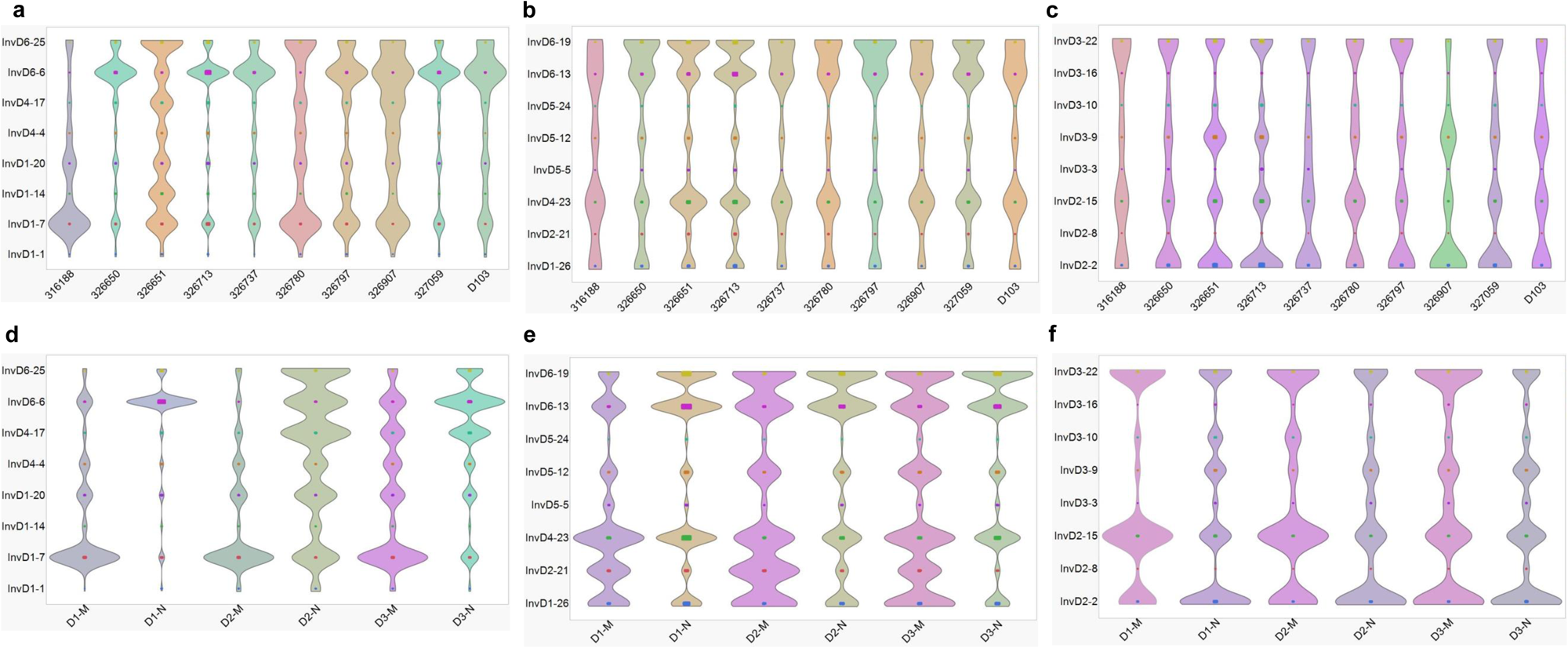
Distribution of InvD gene usage across donors. Comprehensive analysis of the usage of InvD gene families across different donors’ immune repertoires using violin plots. The color-coded X-axis represents individual donors, while the Y-axis indicates the relative usage of each InvD gene family. The existence and prevalence of all unique InvD gene families within **a**, short (11-18 nt), **b**, medium (19-28 nt), and **c**, long (31-37 nt) occurring across 10 donors, except for InvD7-27 due to its minimal length, are shown. **d-e**, Extended analysis includes an additional 3 donors, further categorizing the donors’ antibodies into naïve (N) and memory (M) classifications. Each violin represents the diversity of gene family usage within a donor’s immune repertoire, with wider violins indicating greater prevalence and broader family utilization. Remarkably, memory antibodies exhibit a broader distribution of specific InvD gene families when compared to naïve antibodies. This broader distribution suggests a potential link between specific InvDs and the enhanced immune response capabilities of memory antibodies.

**Fig. S3.**
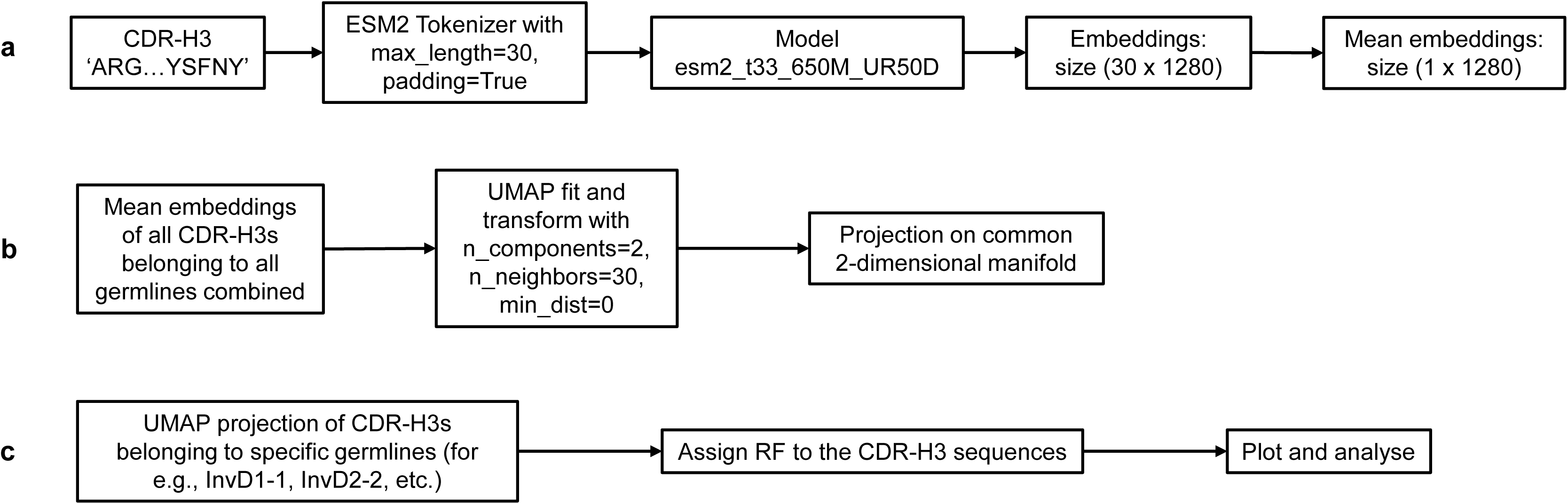
The methodology for zero-shot clustering of InvDs-associated CDR-H3s using mean ESM2 embeddings. **a**, First, CDR-H3 amino acid sequences are tokenized and transformed into vector representations (embeddings) using the ESM2 model (esm2_t33_650M_UR50D variant). **b**, All CDR-H3 sequences are projected onto a common 2-dimensional manifold computed using the UMAP algorithm. **c**, Finally, each InvD sub-families are analysed separately by assigning inverted reading frames (RF1, RF2, and RF3) in the 2-dimensional UMAP manifold.

**Fig. S4.**
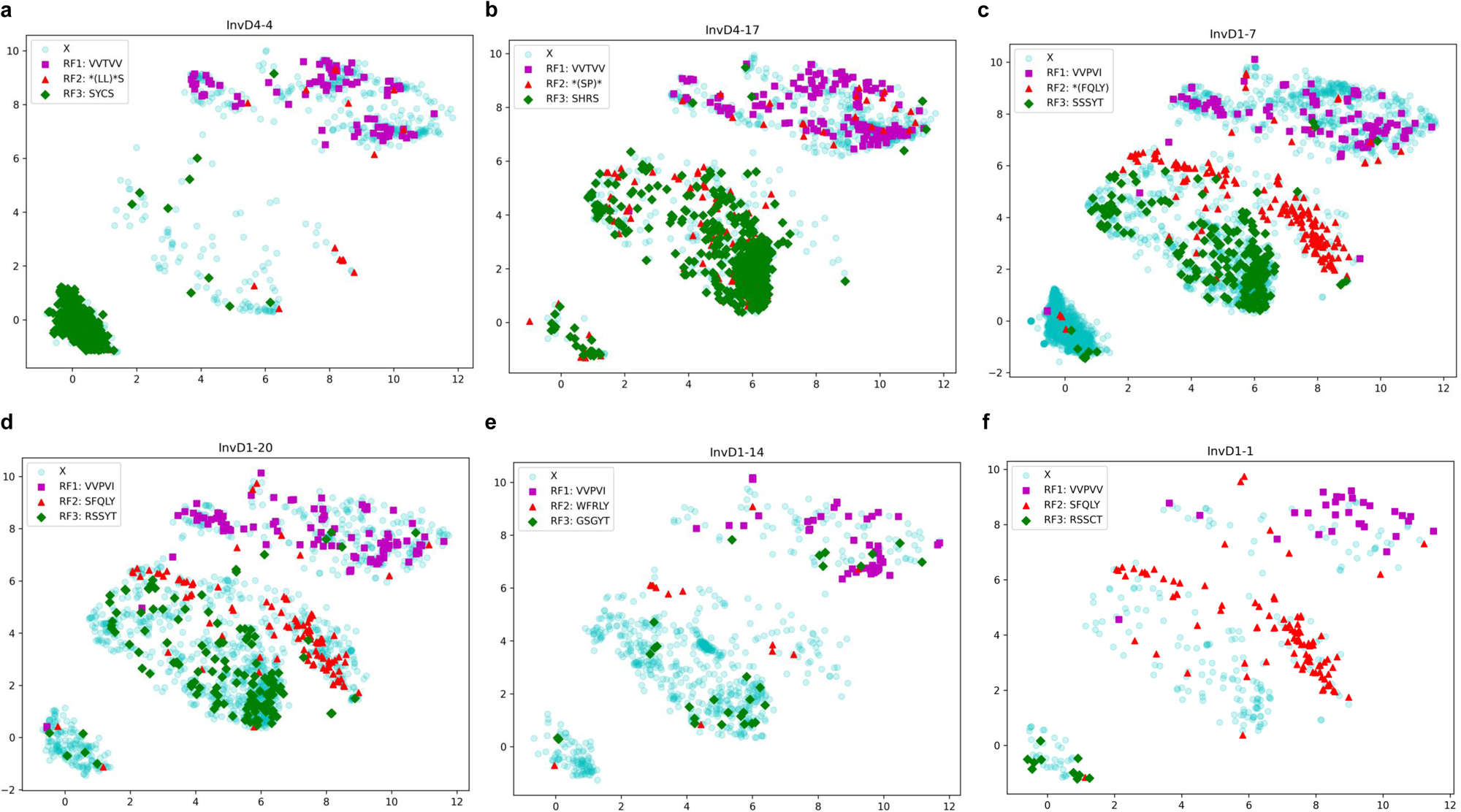
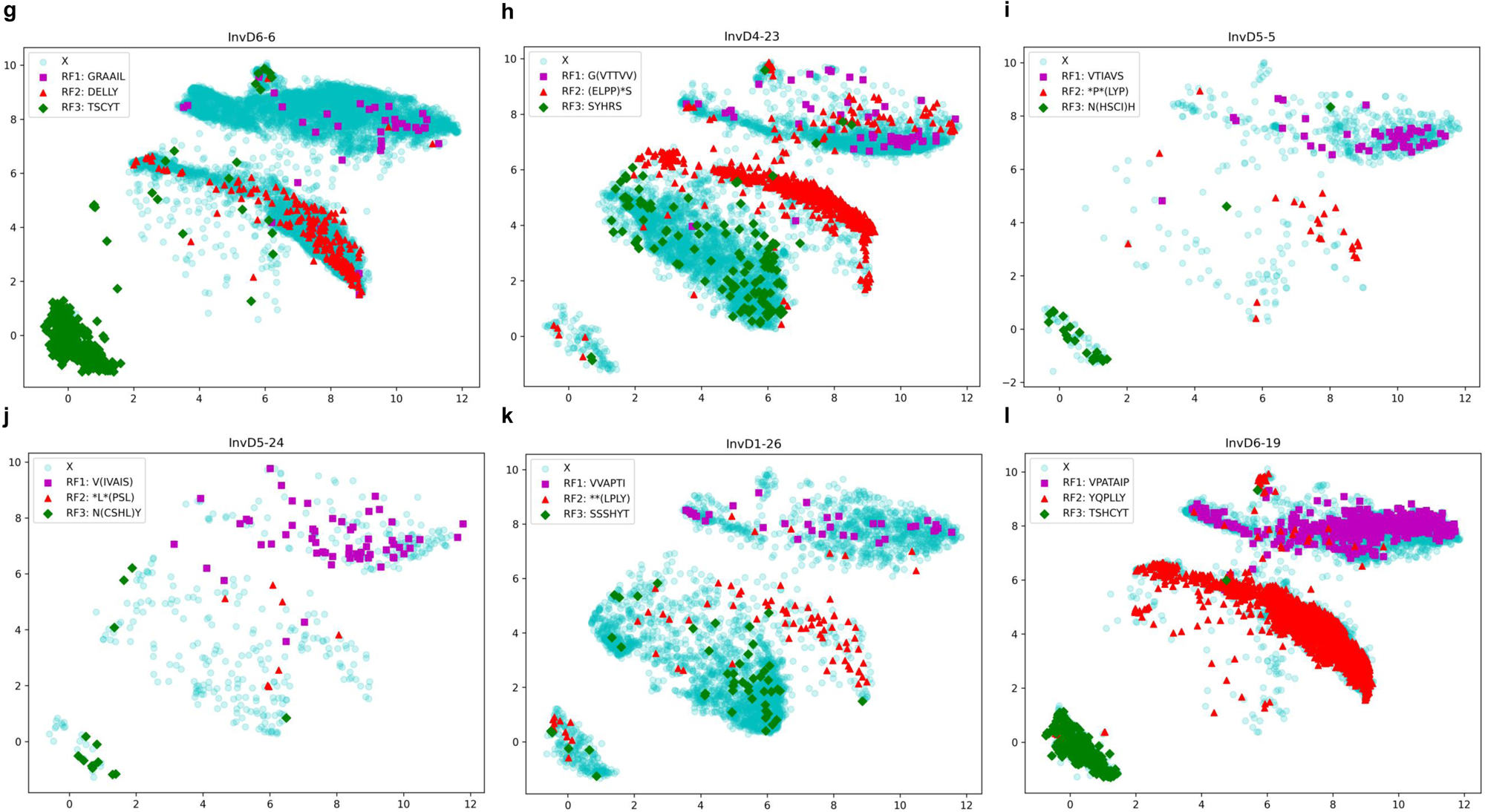
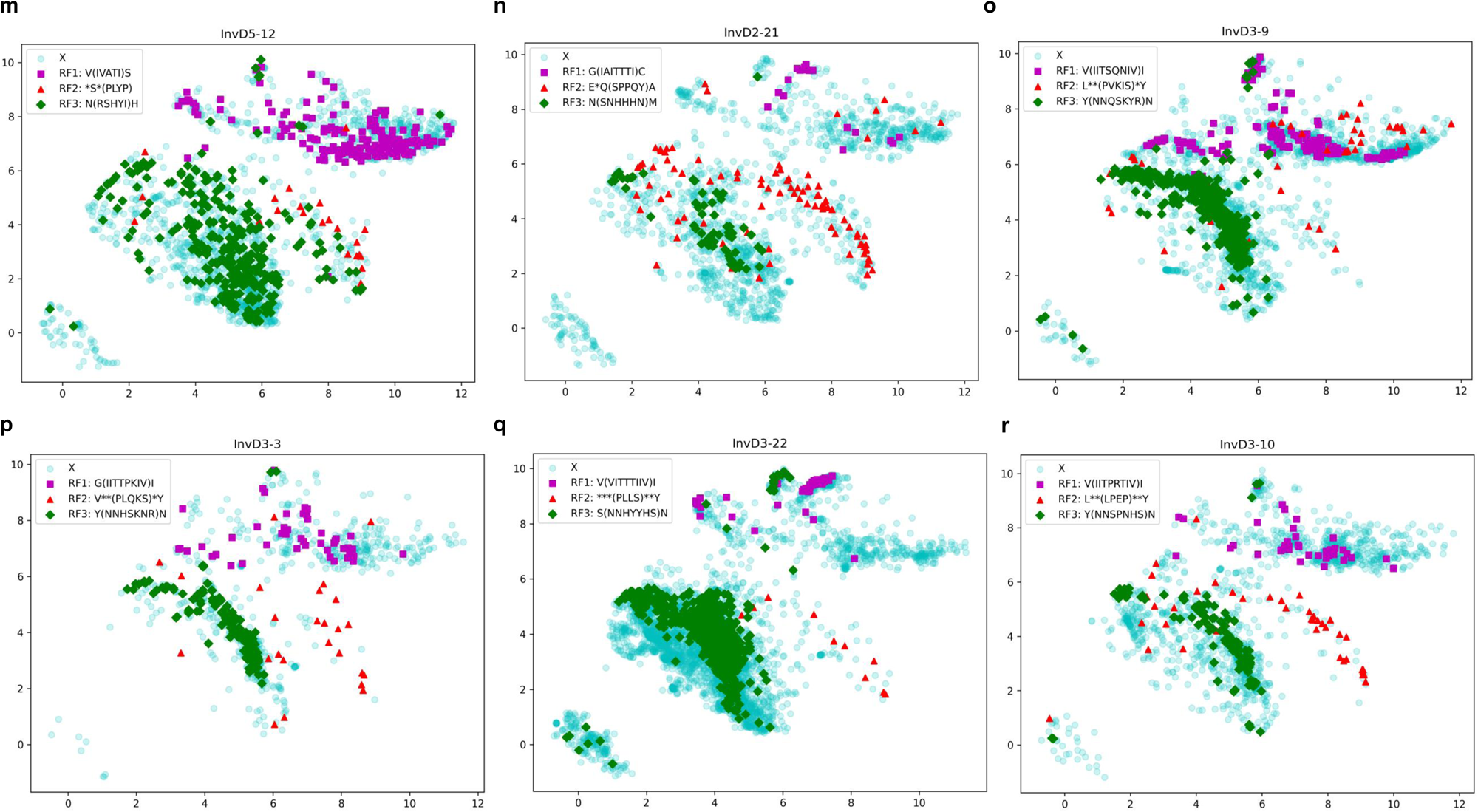
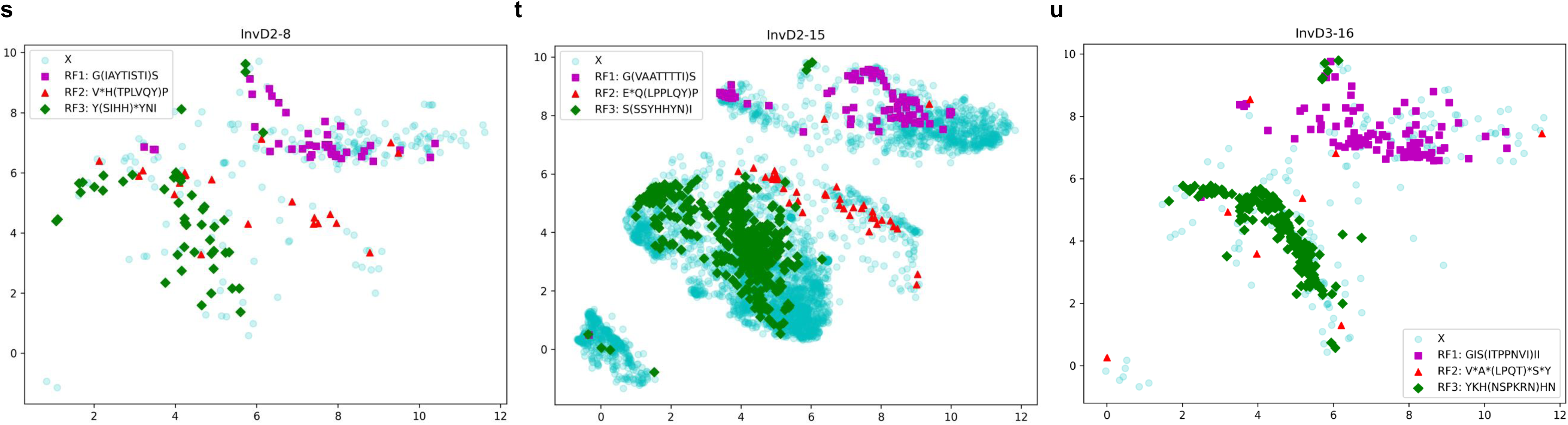
Zero-shot clustering of InvDs-associated CDR-H3s with mapped reading frames (RFs). **a-u**, Clusters of CDR-H3s involving all InvDs, except for D7-27 due to its minimal length, are visualized using the ESM2-based methodology as described in Fig. S3. Inverted reading frames (RF1, RF2, and RF3) are mapped to each InvD-associated CDR-H3 sequence based on a comparison with its germline amino acid sequence. If the full sequence does not match, the motif given within the brackets is used for mapping. The UMAP projections are color-coded according to the assigned reading frames: purple-colored squares for RF1, red-colored triangles for RF2, green-colored diamonds for RF3, and unassigned CDR-H3s are denoted by a cyan-colored circles.

**Fig. S5.**
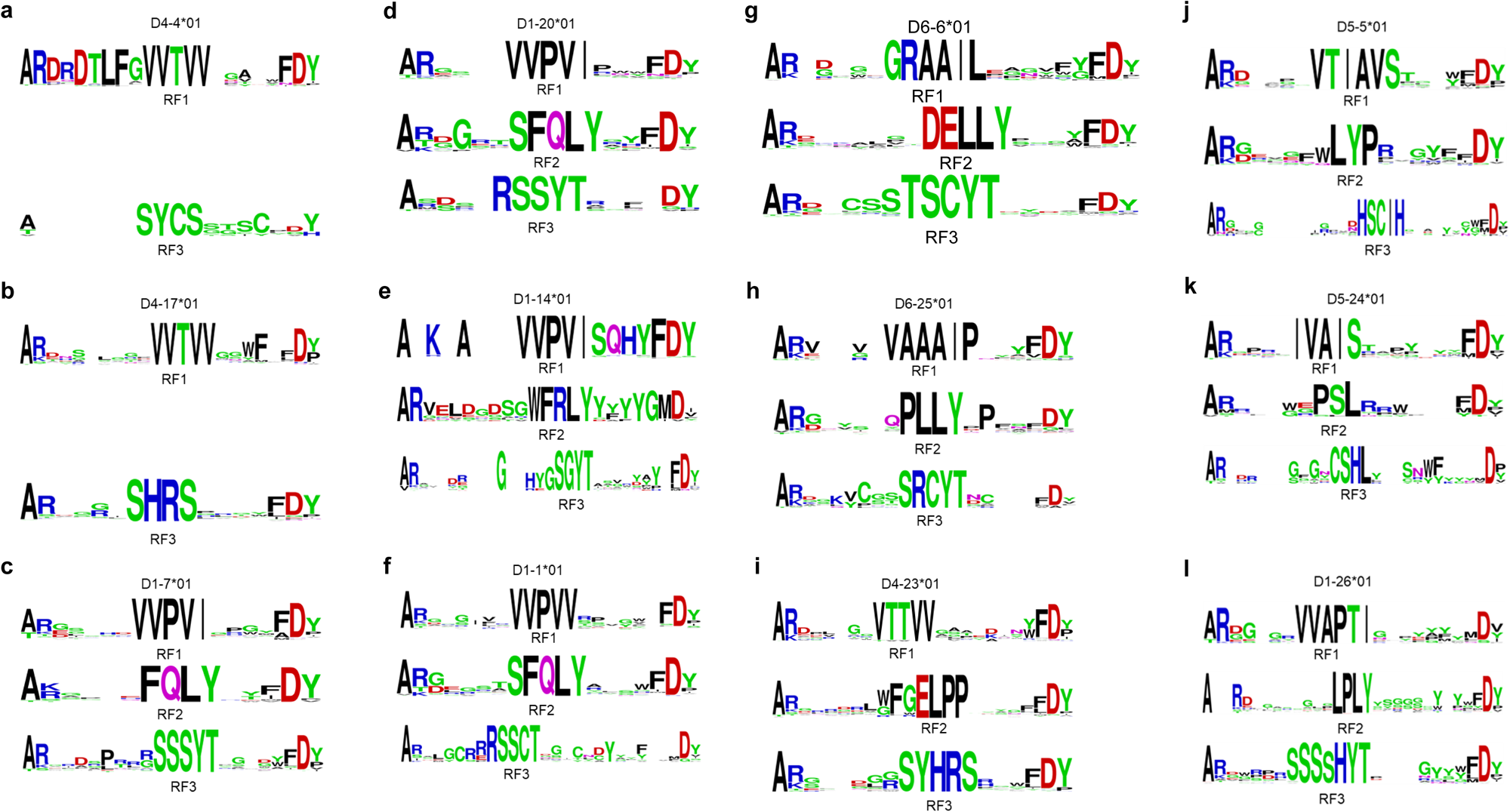
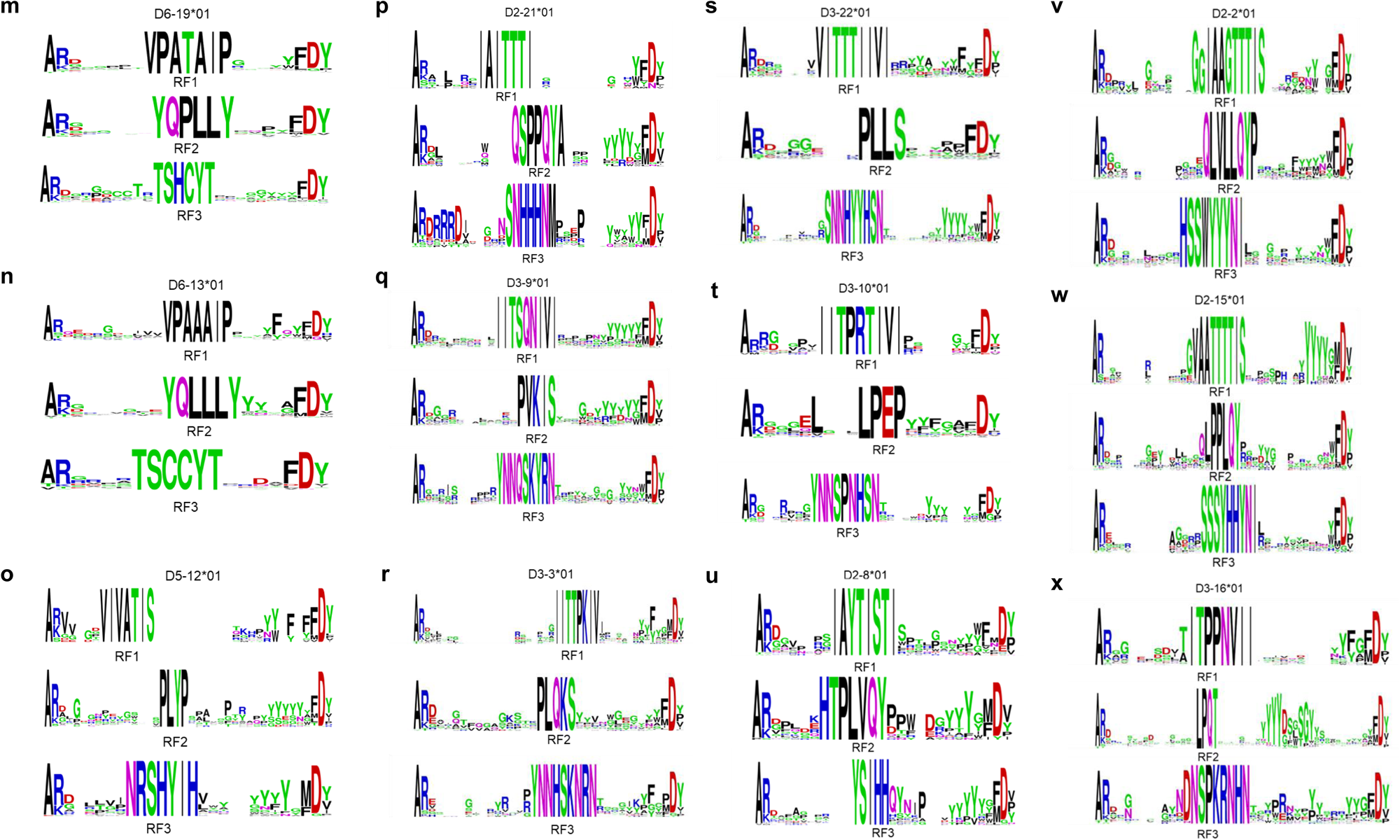
Sequence logos demonstrating the diversity of reading frames (RF1, RF2, and RF3) within germline-encoded InvD-associated CDR-H3s. Sequence logos for all InvDs-associated CDR-H3s, revealing the use of all three reading frames (RF1, RF2, and RF3) in generating diversity within CDR-H3 sequences. Notably, in almost all InvDs, RF2 sequences contain germline-encoded stop codons yet contribute to functional CDR-H3s, suggesting stop codon excision during recombination. Logos for smaller InvD7-27 and RF2s in InvD4-4 and InvD4-17 are excluded.

**Fig. S6.**
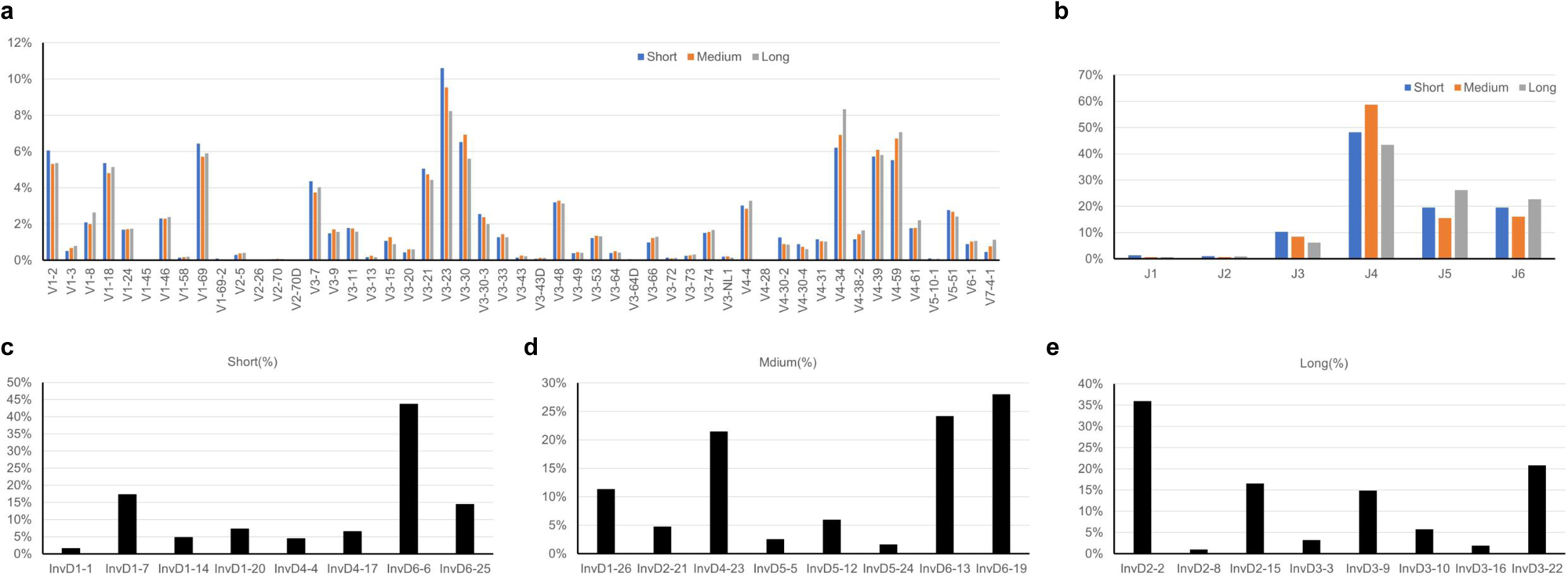
Diversity of V and J gene families in association with InvDs. **a, b**, The patterns of V and J gene family usages across InvD-associated V_H_s are depicted, with no significant variance among short, medium, and long InvDs. **c-e**, Differential frequencies of InvD gene families are shown for short (11-18 nt), medium (19-28 nt), and long (31-37 nt) categories. These findings underscore the diversity and potential functional spectrum of CDR-H3s, generated through V(D)J recombination with InvD involvement within the V_H_s of human antibodies. The X-axis represents gene families, and the Y-axis denotes the percentage of occurrences or frequencies..

**Fig. S7.**
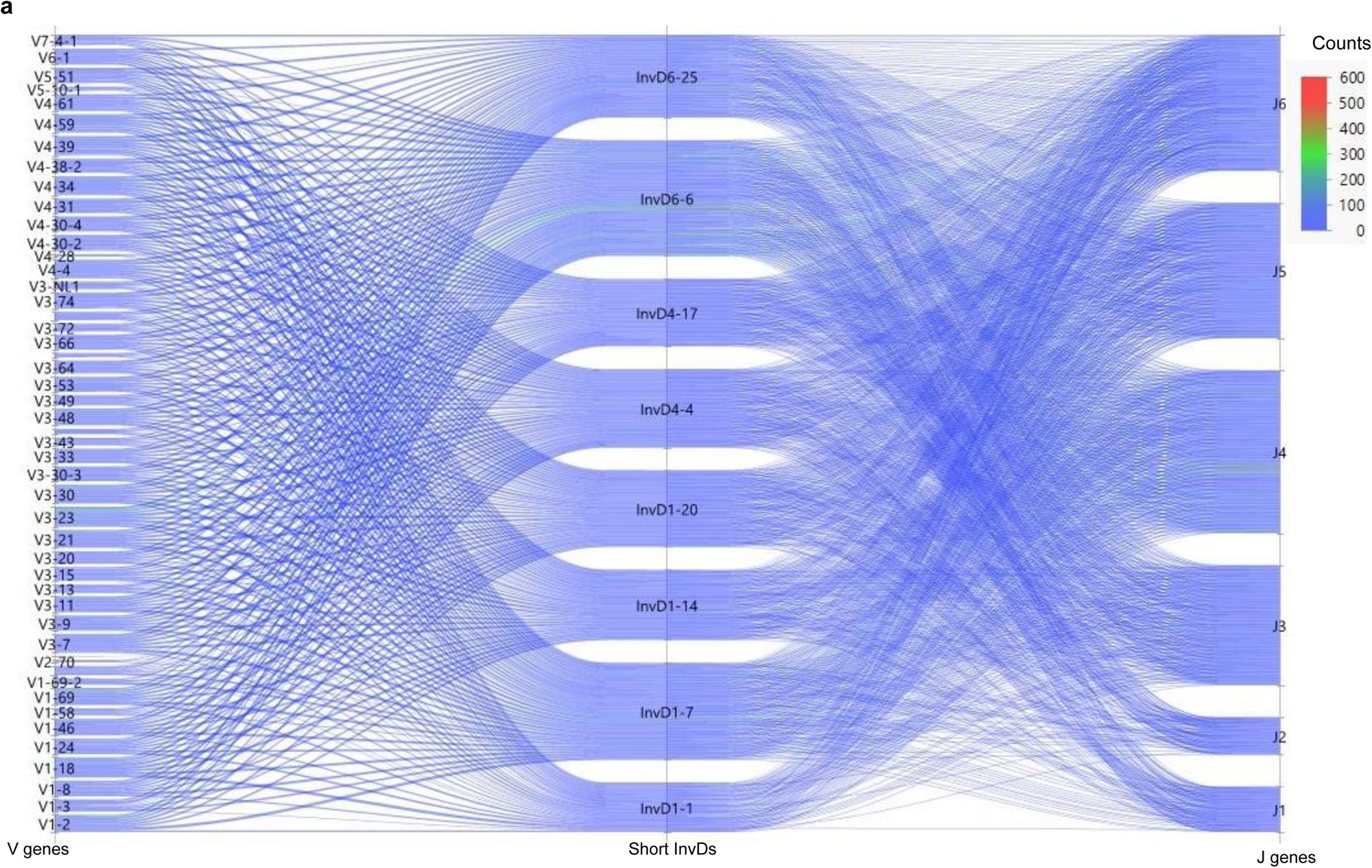
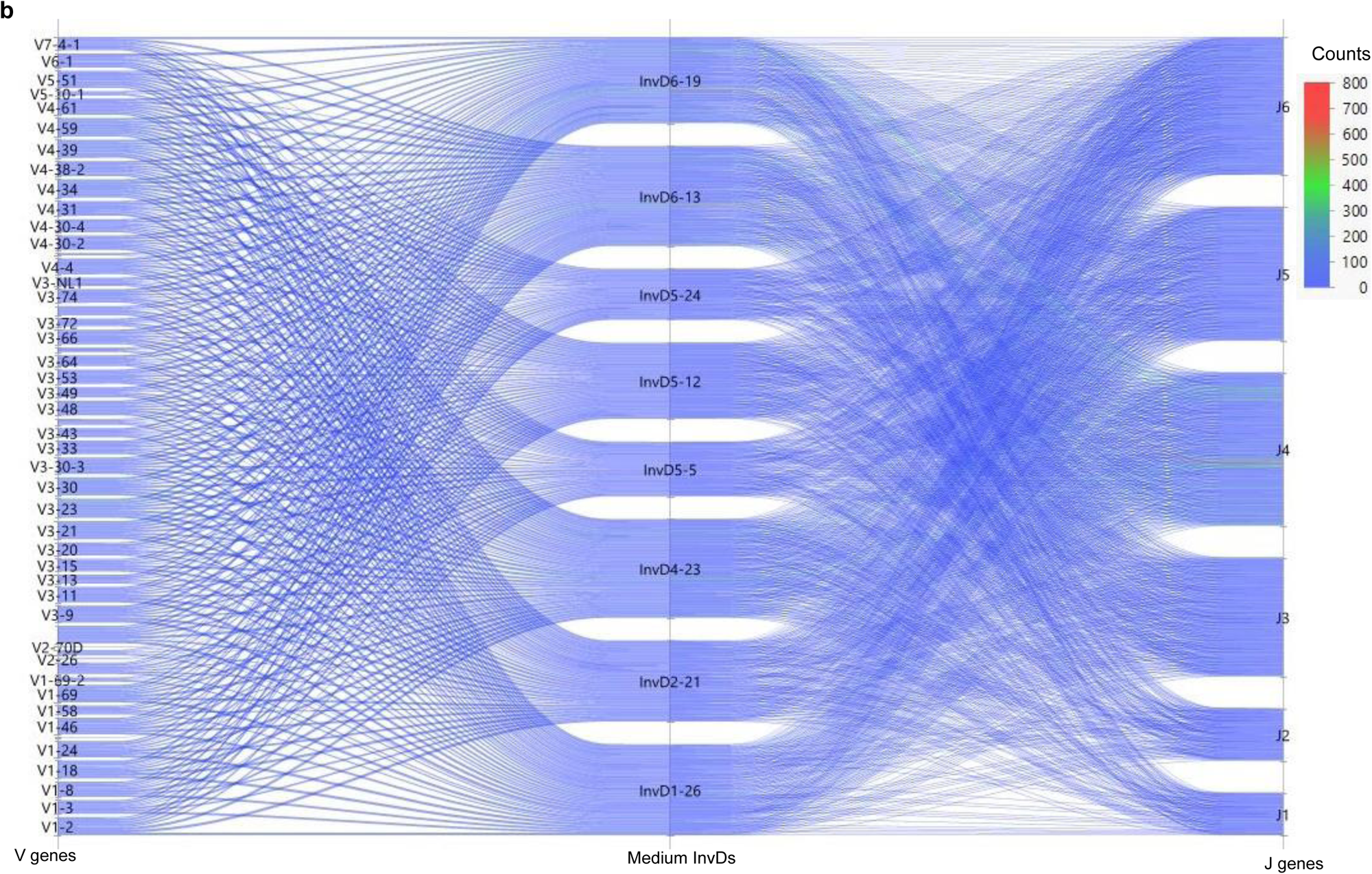
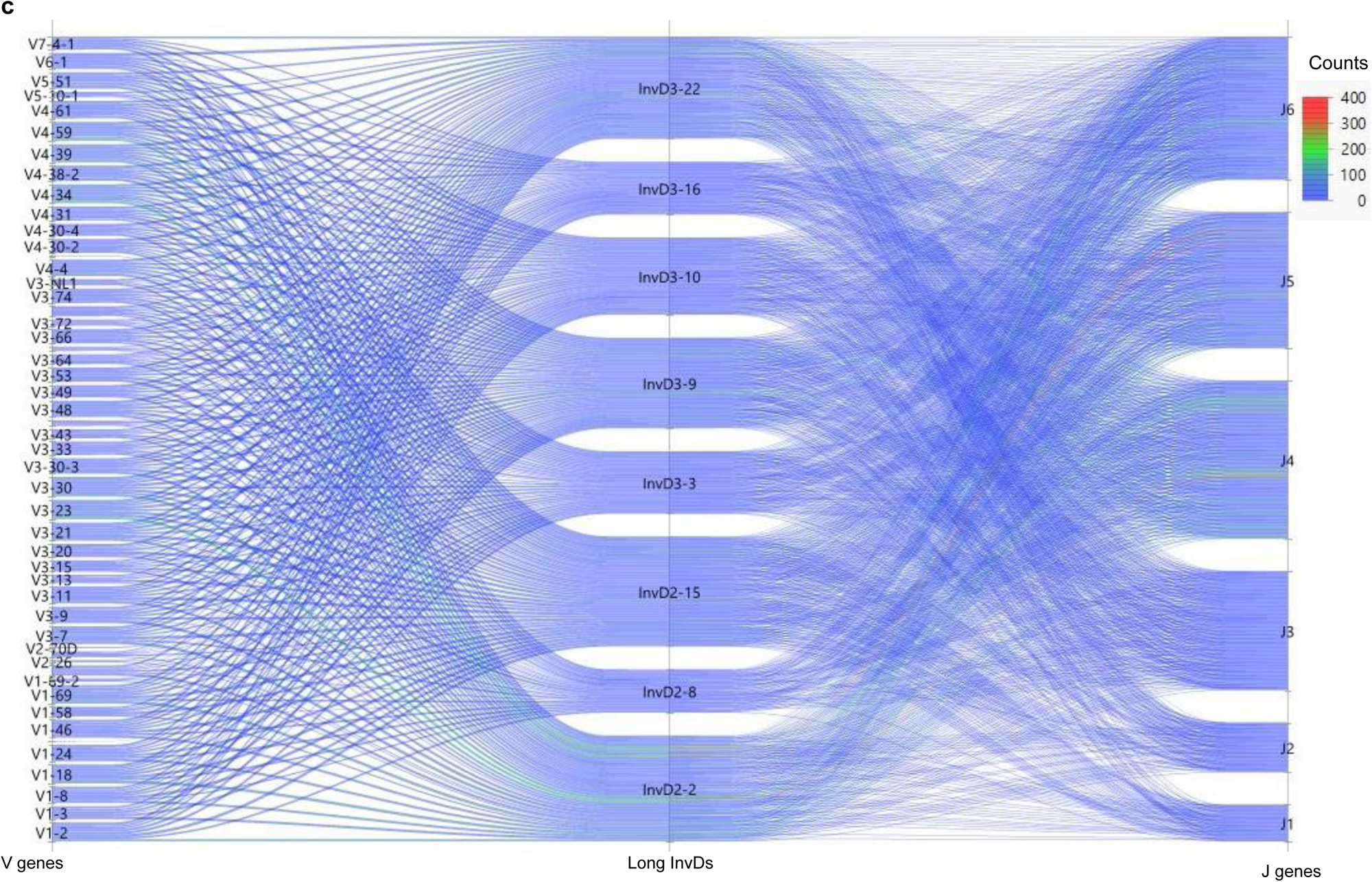
VDJ diversity across InvD categories in human antibodies. VDJ diversity plots illustrate the involvement of InvDs for **a**, short (11-18 nt), **b**, medium (19-28 nt), and **c**, long (31-37 nt) InvDs categories. These plots capture the extensive diversity and potential functionality of V(D)J recombinations with InvD involvement, with a color gradient that signifies the frequency of these recombination events.

**Fig. S8.**
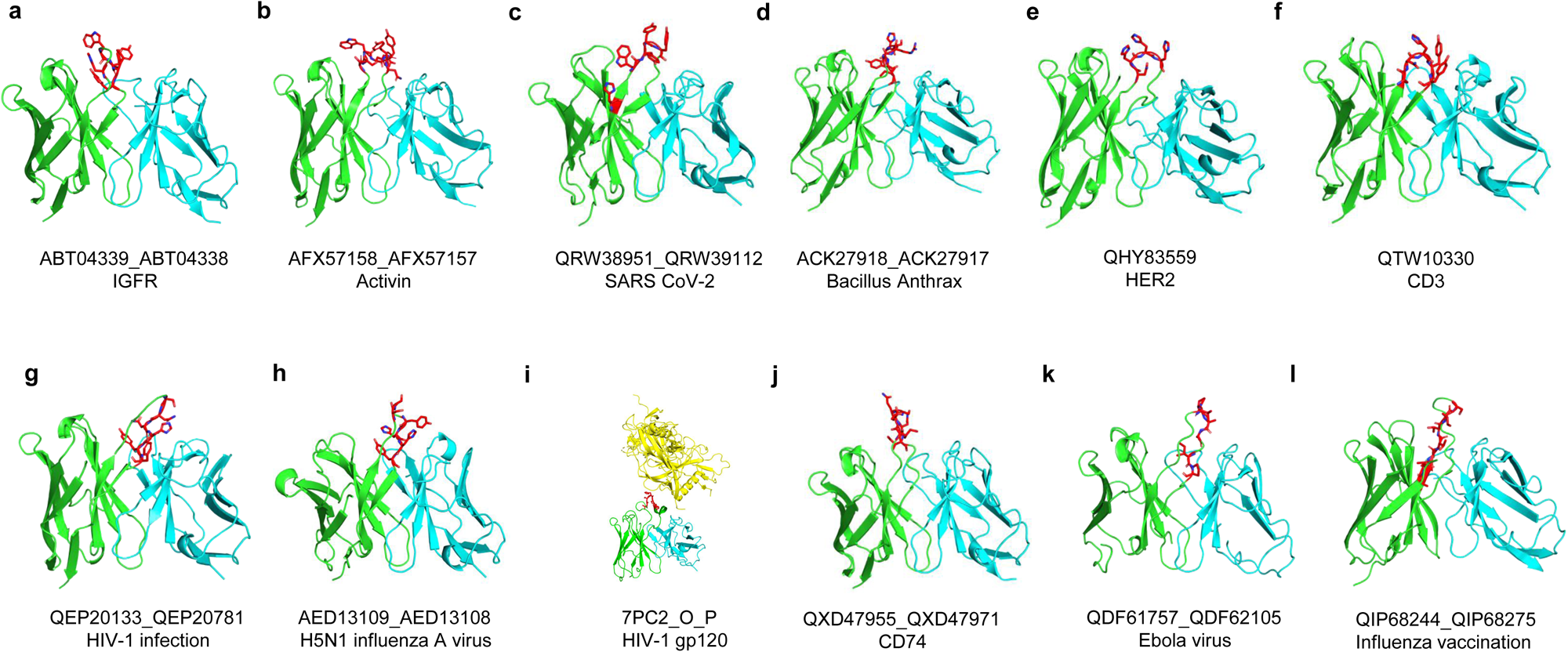
Functional human antibodies harboring InvDs in CDR-H3. **a-l**, Three-dimensional structural models of human antibodies containing InvDs within their CDR-H3s identified in the Patent and Literature Antibody Database (PLAbDab) (details in Supplementary Table 9). The heavy chain is shown in green, and the light chain is shown in cyan, except for panel i, which presents a complex crystal structure (PDB 7PC2) with the antigen in yellow. Germline residues matching the InvDs within the CDR-H3 regions are highlighted in red and displayed as atom-colored stick models using PyMOL software. Additionally, the corresponding PLAbDab IDs and the names of the targets as listed in the database are provided for each antibody model.

**Fig. S9.**
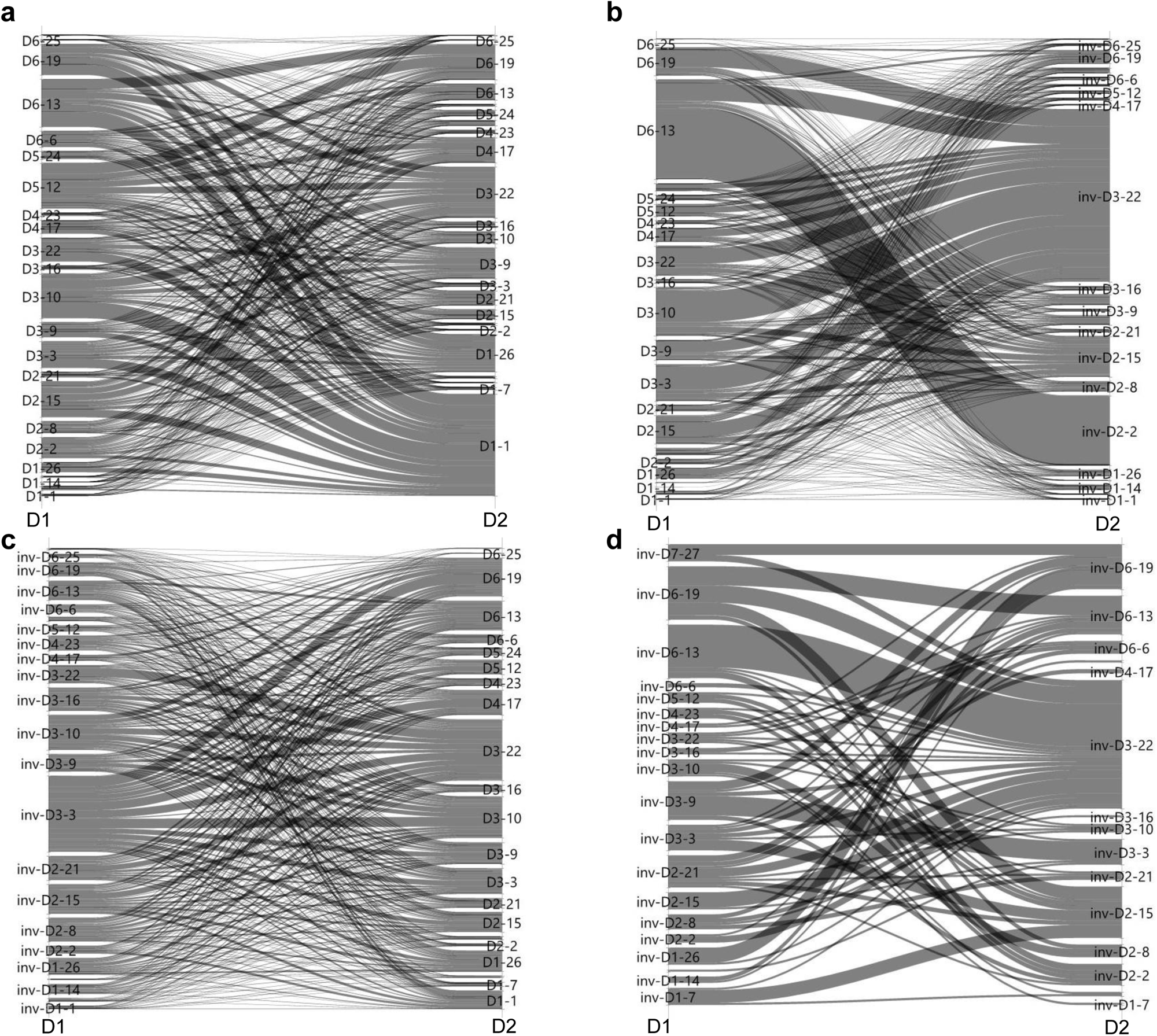
DD fusions involving InvDs create additional diversity in human antibody repertoires. Parallel plots detail the four distinct scenarios involving DD fusions highlighting the intricate diversity within human antibody repertoires through **a**, D-D, **b**, D-InvD, **c**, InvD-D, and **d**, InvD-InvD combinations. The visualized connections underscore the variety of recombinations and the expanded diversity these interactions contribute to the overall antibody repertoire. Line intensities correlate with the occurrence rate of these fusions, mapping a network of potential antibody functionalities emerging from these genetic combinations.

**Fig. S10.**
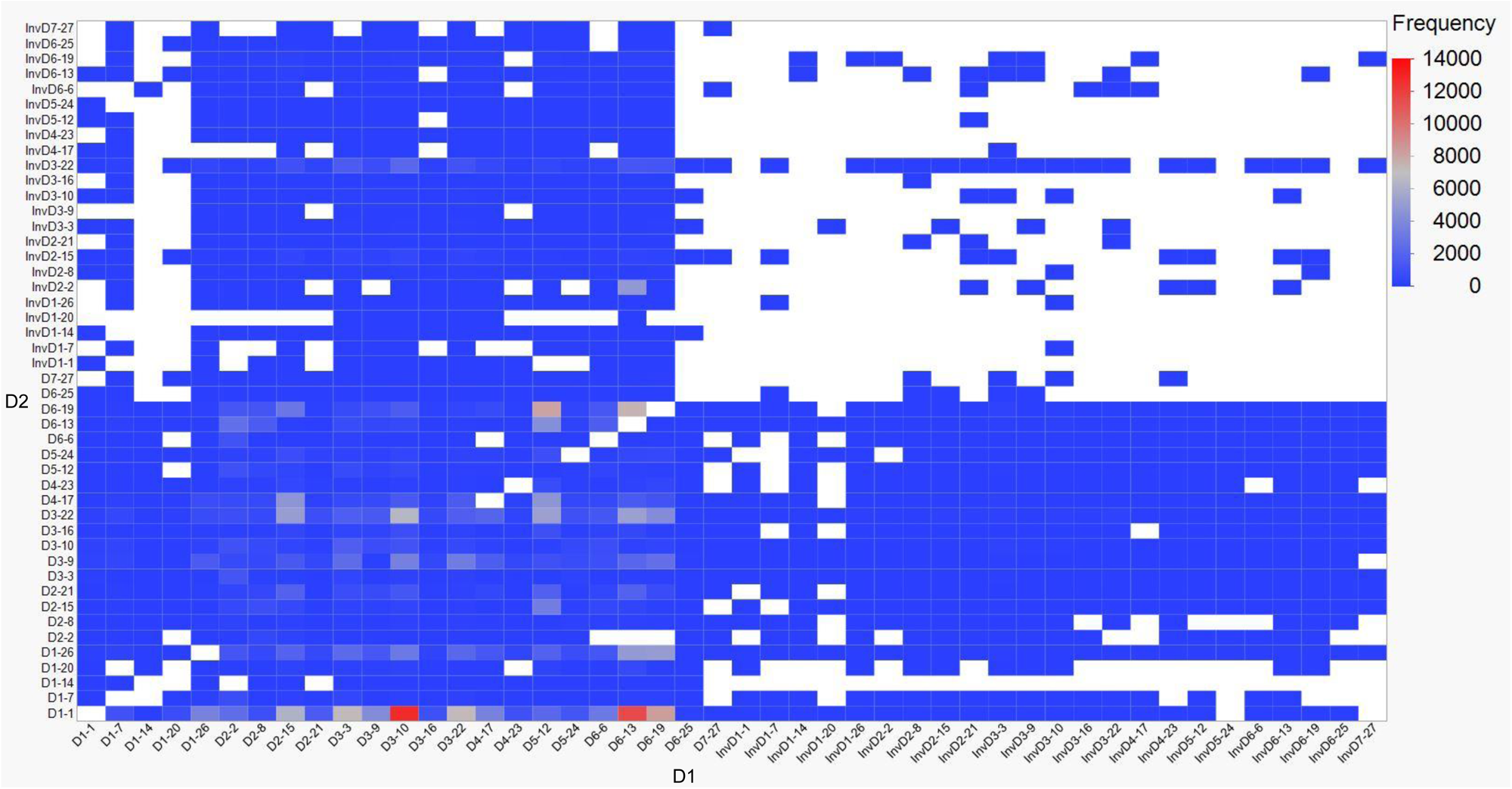
Heat map of DD fusions involving InvDs. The heat map provides a detailed representation of DD fusions in human antibodies, where the X-axis represents the first D gene segment (D1) and the Y-axis represents the second D gene segment (D2) in a tandem orientation. The intensity of the color corresponds to the frequency of fusion events between these segments, capturing the four distinct scenarios: D-D, D-InvD, InvD-D, and InvD-InvD.

## Notes

### Competing Interest Statement

All authors were employees of Sanofi at the time this study was conducted.

